# Towards domain-general predictive coding: Expected TMS excites the motor system less effectively than unexpected stimulation

**DOI:** 10.1101/2020.02.18.953562

**Authors:** Dominic M. D. Tran, Nicolas A. McNair, Justin A. Harris, Evan J. Livesey

## Abstract

The brain’s response to sensory input is modulated by prediction. For example, sounds that are produced by one’s own actions, or those that are strongly predicted by environmental cues, are perceived as less salient and elicit an attenuated N1 component in the auditory evoked potential. Here we examined whether the neural response to direct stimulation of the brain is attenuated by prediction in a similar manner. Transcranial magnetic stimulation (TMS) applied over primary motor cortex can be used to gauge the excitability of the motor system. Motor-evoked potentials (MEPs), elicited by TMS and measured in peripheral muscles, are larger when actions are being prepared and smaller when actions are voluntarily suppressed. We tested whether the amplitude of MEPs was attenuated under circumstances where the TMS pulse can be reliably predicted, even though control of the relevant motor effector was never required. Self-initiation of the TMS pulse and reliable cuing of the TMS pulse both attenuated MEP amplitudes, compared to MEPs generated programmatically in an unpredictable manner. These results suggest that predictive coding may be governed by domain-general mechanisms responsible for all forms predictive learning.

## Main Text

Making predictions about future events is one of the most critical and dynamic functions of the human brain. Converging evidence suggests that the human brain forms predictions based on mental models of the world, which are updated based on the outcomes that we subsequently experience (Bar, 2007; Schultz & Dickinson, 2000). Predictive learning of this nature allows us to anticipate changes in our environment and adjust our behaviour to enhance good outcomes and avoid undesirable ones. While our deliberate actions clearly benefit from learning to explicitly predict outcomes, evidence suggests that predictive learning extends far deeper, with the brain automatically generating predictions at every level of information processing from early perception to reward coding, and motor control (Chong, Familiar, & Shim, 2016; Petro & Muckli, 2016; Schultz, Dayan, & Montague, 1997; Wacongne et al., 2011; Yeung, Botvinick, Cohen, 2004). These predictions affect the way external events are ultimately perceived and interpreted, allowing the brain to distinguish between novel or important information and irrelevant or redundant information. For instance, sensory stimulation tends to be perceived as less salient when the sensation arises from a self-initiated action, such as speaking, tapping, or interacting with an object. Take the sound of footsteps as a specific case: the sound in a deserted street late at night does not raise alarm if it is generated by one’s own footsteps, but the same sound becomes highly salient if the movements and sounds do not align. Walking is highly predictive of hearing footsteps, thus the action itself provides a source of internal prediction for the auditory sensations (Crapse & Sommers, 2008b; Poulet & Hedwig, 2007; Whitford et al., 2011). It has been suggested that this form of sensory attenuation to stimulation produced by one’s own actions is the reason we are unable to tickle ourselves (Blakemore, Frith, & Wolpert, 1999; Blakemore, Wolpert, & Frith, 2000; Weiskrantz, Elliott, & Darlington, 1971).

The sensations produced by our own actions are not only subjectively less salient, but the neural signals associated with processing sensory information are objectively attenuated. The onset of sensory input, for example an auditory tone, triggers a cascade of neural activity that is measurable as an event-related potential (ERP) using encephalography (EEG). An early component of the auditory ERP, the N1, varies in amplitude with the loudness of the tone (Mulert et al., 2005; Simmons, Nathan, Berger, & Allen, 2011). Critically, a self-generated tone (triggered via a button press) produces a smaller N1 component compared with the same tone when played spontaneously (e.g., Elijah et al., 2016). This suppression of N1 is also accompanied by a reduction in the perceived loudness of the tone. This effect has been interpreted as a consequence of a *corollary discharge*: an internal signal generated when performing an action, which modulates sensory processing by routing copies of the movement command to inform the sensory system of an upcoming action (Blakemore, Frith, & Wolpert, 1999). Corollary discharge has been documented in different sensory systems of both invertebrates and vertebrates (Crapse & Sommer, 2008a), and is thought to help organisms prioritise processing sensory inputs generated from the external world, which tend to be more biologically relevant than self-generated sensation. However, similar sensory attenuation effects are also produced when an externally-generated tone is made highly predictable by other external events, such as a preceding warning cue (Ford et al., 2007). These results suggest that it is the prediction or expectation of sensory input, more generally, that changes how the sensation is ultimately processed by the brain.

One way prediction can modulate the perceptual and neural salience of sensory events is through learning mechanisms that continually update based on the discrepancies between anticipated and experienced events. These mechanisms are vital for us to learn about new, unanticipated environmental changes. The neural representation of predictions and prediction error (i.e., predictive coding) is considered to be a fundamental, ubiquitous, and highly adaptive property of the healthy brain (Clark, 2013). Moreover, psychopathologies related to psychosis, particularly schizophrenia (Ford et al., 2001; Horga et al., 2014; Shergill, Samson, Bays, Frith, & Wolpert, 2005), as well as autism (Van Boxtel & Lu, 2013; Van de Cruys et al., 2014), are associated with deficits in both predictive coding at a perceptual level and explicit predictions based on reasoned beliefs. Recent evidence of the functional similarities in predictive learning at different stages of information processing, from higher-order conscious predictions about the causal relationships between events to automatic processes such as sensory attenuation, has transformed contemporary theories in psychology and cognitive neuroscience (Miller & Cohen, 2001; Sterzer et al., 2018) and led some to argue that prediction error may be the *lingua franca* of the brain (Bastos et al., 2012; Friston & Kiebel, 2009).

The research on sensory attenuation naturally examines changes in sensory processes based on environmental events with which we have had a rich developmental experience and long evolutionary history (e.g., sights, sounds, and touch; Shergill, Bays, Frith, Wolpert, 2003). That is, we have evolved specific sensory mechanisms and fine-tuned their function with years of personal experience. What about neural events with which we have limited experience? Advances in direct neural stimulation offer the chance to ask whether similar prediction effects observed in well-experienced and evolutionarily-refined systems such as vision and audition, are present across other neural systems. If predictive coding mechanisms are domain-general and exist across different neural systems, we would expect to see similar prediction effects beyond the sensory system.

In recent years, a wealth of innovative research has emerged that investigates corticospinal excitability using transcranial magnetic stimulation (TMS). TMS is a form of non-invasive brain stimulation in which cortical neurons are excited by a coil placed on the scalp. TMS applied over the primary motor cortex (M1) can be used to study the function of the motor system by measuring responses to the stimulation in peripheral muscles controlled by M1 via the pyramidal tracts. These responses are quantified by measuring motor-evoked potentials (MEPs) with electromyography (EMG). The peak-to-peak amplitude of the MEP provides an index of corticospinal excitability at the time of stimulation and is sensitive to voluntary action preparation and suppression (Poole et al., 2018), as well as automatic effects triggered by stimuli that are linked to specific actions (Seet et al., 2019; Tran et al., 2019; Tran et al., online ahead of print). Converging evidence thus suggests that motor excitability, as indexed by MEP amplitude, is sensitive to predicting the need to act (response preparation) or predicting the need to suppress a response (proactive response inhibition).

A question that has not been asked is whether the sensitivity of the motor cortex to TMS is also susceptible to prediction more generally (even in the absence of any response requirements). That is, can the brain quickly learn to anticipate direct neural stimulation and, if so, is its response to this stimulation attenuated accordingly? Here we used TMS to investigate this question by stimulating M1 with TMS when the contralateral hand is completely at rest, but under conditions that manipulate the predictability of the TMS pulse itself. In doing so, we hope to test whether direct stimulation of the motor system reveals similar predictive coding properties observed in the sensory systems and other predictive learning contexts.

Based on the sensory attenuation literature, we hypothesise that self-generated or predictable TMS pulses will result in reduced corticospinal excitability, as indexed by MEP amplitude, compared to TMS pulses triggered in an unpredictable manner. While attenuation effects may be unique to the functional architecture of the sensory cortex— possibly evolved to serve a specific role because of the close and consistent relationship between actions and their perceptual consequences—some evidence suggests that these effects are modifiable with training (Elijah et al., 2018). That is, experience may play an important role in shaping the underlying predictive coding mechanisms that modulate the way in which the brain responds to external events. Further, assuming the general function of attenuation effects is to modulate how we are affected by predicted and unpredicted external events, we expected that prediction modulation should also influence the brain’s response to novel, direct stimulation of the motor system.

## Results

### Experiment 1

Experiment 1 assessed the sensitivity of the motor system to self-generated (predictable) and computer-generated (unpredictable) stimulation using single pulse TMS. By systematically varying the intensity of TMS pulses delivered to left M1, we mapped the stimulus-response (S-R) function of MEP amplitudes recorded from the right first dorsal interosseous (FDI) muscle. Separate curves were fitted for pulses generated by a button press using the left thumb or left foot as well as pulses generated programmatically. The left foot was included as it is possible that self-generation of the TMS pulse using the hand ipsilateral to the stimulated motor cortex may trigger inhibitory or excitatory changes in MEPs simply due to direct inter-hemispheric communication between the motor cortices. However, Labruna et al. (2014) have demonstrated that while MEPs in the left FDI muscle are indeed suppressed during preparation of a motor response involving a non-homologous muscle in the opposite hand, this suppression disappears when preparing a contralateral motor response with a non-homologous effector such as the foot.

**Figure 1.**
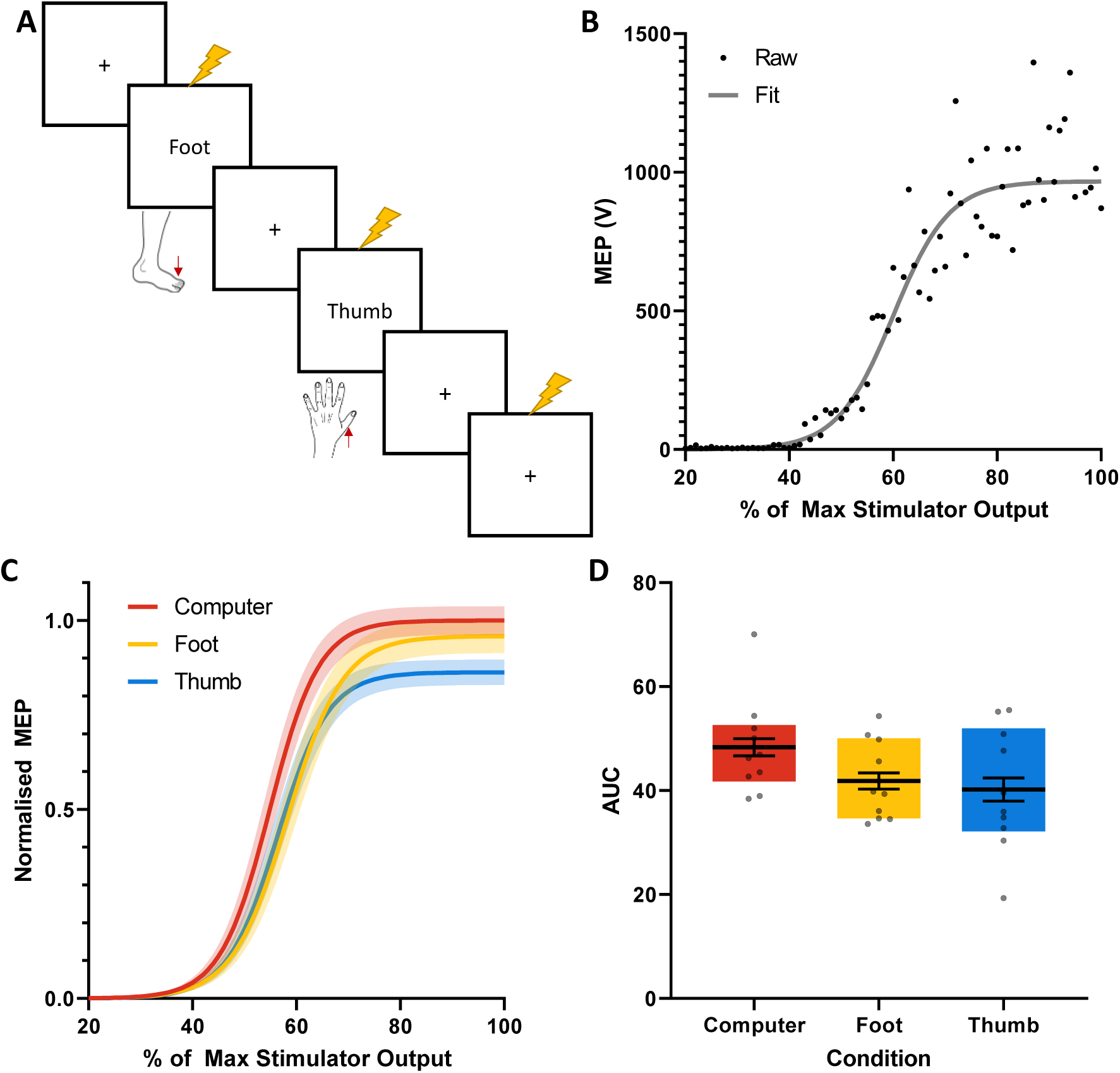
Schematic representation and results of Experiment 1. **A)** Squares represent the screen displayed to participants. A fixation cross (**+**) was used as the default presentation. The word “Foot” indicated that participants should press the spacebar with their left foot when ready (on a keyboard positioned the under the table). The word “Thumb” indicated that participants should press the spacebar with their left thumb when ready (on a different keyboard positioned on the table). TMS was triggered immediately on the press of the spacebar for both self-generated conditions. TMS was also randomly triggered automatically during the fixation cross (Computer condition). **B)** Raw MEP data and sigmoid fit from one condition for an example participant. **C)** S-R curves mapping MEP amplitude (normalised to the Computer condition response for each participant) to stimulator intensity. **D)** Area under the S-R curve for each condition. Black bars represent means, and error bars represent within-participant standard errors (Morey, 2008). Dots represent individual data points, and filled colour boxes represent interquartile range.

Mean goodness of fit and parameter values for sigmoid functions are reported in the Supplementary Materials. The S-R curves for self-generated TMS pulses, triggered using either left thumb or their left foot, had midpoints at higher stimulator outputs (abscissa) and lower maximum MEP amplitudes (ordinate) compared to the S-R curve for computer-generated TMS pulses. The area under the curve for computer-generated pulses was significantly larger than that of the self-generated pulses from both the thumb (*t*(9) = 2.77, *p* = 0.022, *d* = 0.88) and foot (*t*(9) = 3.43, *p* = 0.007, *d* = 1.09). The thumb- and foot-generated curves did not differ significantly from each other (*t*(9) = 0.58, *p* = 0.711, *d* = 0.18). Given the two informal outliers (not > 3 standard deviations beyond the mean) in the computer and thumb conditions, we also analysed the data using a set of non-parametric Wilcoxon signed-ranked tests and found a statistically consistent pattern of results: computer > thumb (*W* = 41, *p* = 0.037), computer > foot (*W* = 49, *p* < 0.010), thumb vs foot (*W* = 9, *p* = 0.695). These results provide a proof-of-concept demonstration that the predictability of TMS pulses modulates the amplitude of the resulting MEP; with corticospinal excitability suppressed for self-generated (i.e., predictable) TMS pulses compared to unpredictable pulses.

### Experiment 2

Experiment 2 teased apart two predictive components of the self-generated TMS conditions — the visual cue and the action itself — to examine whether an action (i.e., a key press) was necessary for generating the effects found in Experiment 1. Although the actions to initiate TMS in Experiment 1 were voluntary and self-paced, the visual prompt still provides predictive information about the imminent delivery of a TMS pulse. The attenuation observed in Experiment 1 may therefore result from a combination of prediction effects based on visual and self-initiation cues. To test this, we used the same conditions as Experiment 1 (self-generated with left thumb or foot, and unpredictable computer-generated), but also included a predictable computer-generated condition in which a warning cue signalled the onset of a TMS pulse. Pulse intensity was fixed to 120% of the resting motor threshold (rMT) for each participant. For the self-generated condition, half the participants triggered the TMS pulse using their left thumb, and the other half triggered the TMS pulse using their left foot. For the computer-generated conditions, the TMS pulse was programmatically triggered either unpredictably or following the presentation of a visual stimulus (warning cue). For the predictable computer-generated condition, the timing of the TMS pulse relative to the warning cue onset was yoked to the participant’s response time on the previous self-generated trial, so that the timing of the TMS pulse relative to the visual cue was identical across the self-generated and warning cue conditions. Response time to the visual cue varied very little within participants (Supplementary Materials). Experiment 2 also investigated the effect of prediction contingency on corticospinal excitability by having two distinct phases: one where the contingency of pressing the TMS trigger key (self-generation condition) and the presentation of the warning cue resulted in 1 TMS pulse pseudorandomised in every 4 trials (low contingency condition); and a second phase where every key press and warning cue presentation resulted in a TMS pulse (high contingency condition). We expect attenuation to be stronger when the predictive relationships between the cues and the TMS are stronger (i.e. under higher contingency).

**Figure 2.**
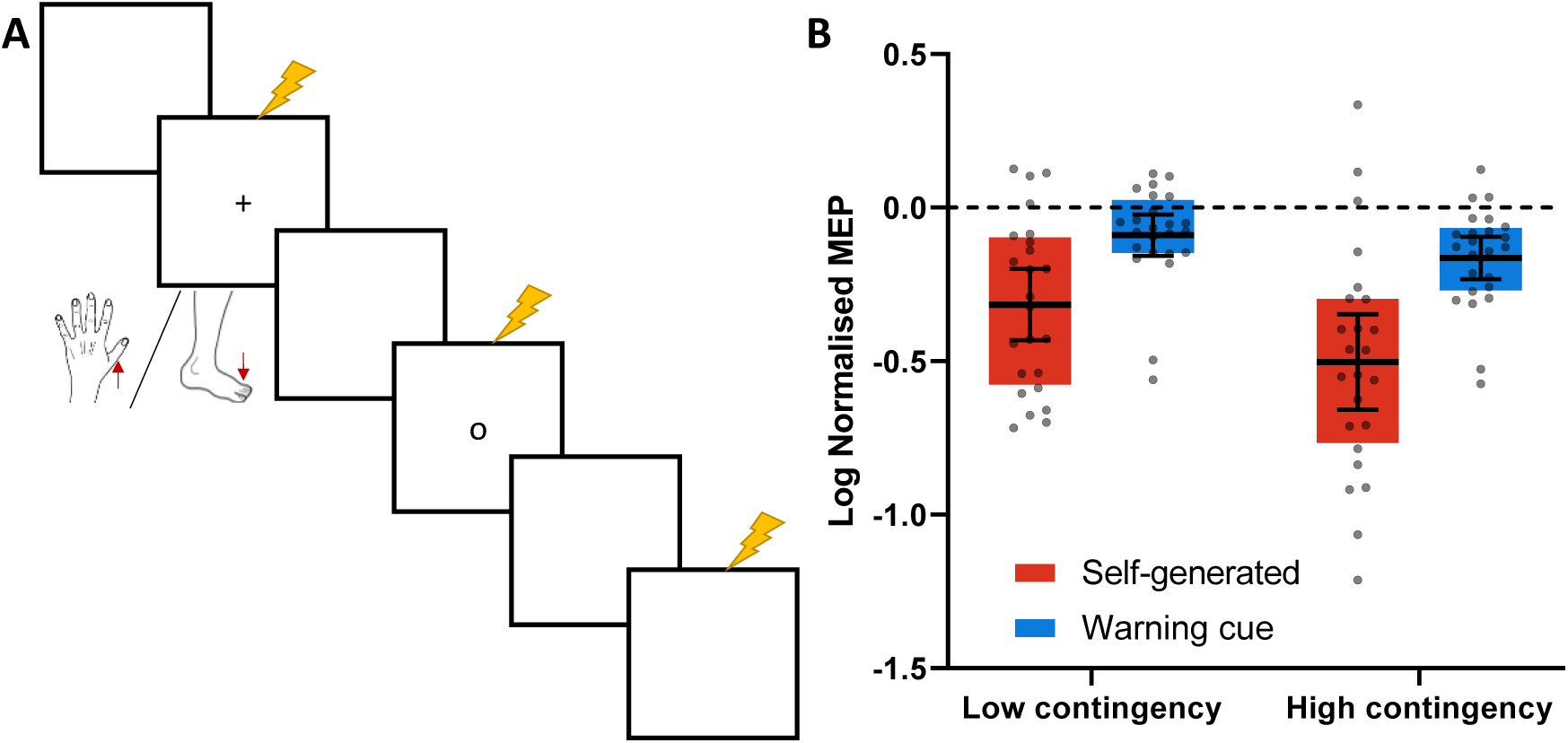
Schematic representation and results of Experiment 2. **A)** Squares represent the screen displayed to participants. A blank screen was used as the default presentation. Participants were instructed at the beginning of the experiment that one shape (e.g., “**+**”) signalled that they should press the spacebar when ready (Self-generated condition) while another shape (e.g., “**o**”) signalled that they did not need to make a response (Warning cue condition). TMS was triggered immediately on the press of the spacebar for the Self-generated condition and after a short delay (see text for details on the timing) for the Warning cue condition. TMS was also randomly triggered during the blank screen (Unpredictable condition). **B)** Log MEPs normalised to the Unpredictable condition for each participant. Black bars represent means, and error bars represent 95% CI. Dots represent individual data points, and filled colour boxes represent interquartile ranges. Dashed line represents baseline of log normalisation.

MEPs from each of the predictable conditions were log normalised to the unpredictable computer-generated TMS pulses (see Method and Materials – Data Analysis for more details). Therefore, deviations from the baseline of 0 indicates a difference between the predictable and unpredictable conditions. Self-generated pulses triggered by the thumb or foot (pooled) showed significantly reduced MEPs during both the low (one-sample t-test H_0_ = 0: *t*(23) = 5.63, *p* < 0.001, *d* = 1.15), and high (one-sample t-test H_0_ = 0: *t*(23) = 6.68, *p* < 0.001, d = 3.07) contingency phases. Similarly, stimulation of the motor cortex by predictable computer-generated TMS pulses (signalled by a warning cue) showed significantly reduced MEPs during both the low (*t*(23) = 2.76, *p* = 0.011, *d* = 0.56) and high (*t*(23) = 4.89, *p* < 0.001, *d* = 1.00) contingency phases. The data were also analysed as a full mixed model ANOVA with effector as a between group factor [thumb, foot], cue type as a within group factor [self-generated, signalled computer-generated], and phase as a within group factor [low contingency, high contingency] to assess the effect of prediction contingency. There were no significant 3-way (*F*(1,22) = 0.26, *p* = 0.614, *η*_*p*_^2^ = 0.012) or 2-way interactions with effector (effector × cue type: *F*(1,22) < 0.01, *p* = 0.954, *η*_*p*_^2^ < 0.001; effector × phase: *F*(1,22) = 0.86, *p* = 0.364, *η*_*p*_^2^= 0.038), nor a main effect of effector (*F*(1,22) = 1.00, *p* = 0.328, *η*_*p*_ ^2^= 0.04). There was a significant main effect of cue type (*F*(1,22) = 26.56, *p* < 0.001, *η*_*p*_^2^ = 0.55), a significant main effect of phase (*F*(1,22) = 11.01, *p* = 0.003, *η*_*p*_^2^ = 0.33), and no significant effector × phase interaction (*F*(1,22) = 2.29, *p* = 0.144, *η*_*p*_^2^ = 0.09).

The results indicate that both self-generated pulses and predictable computer-generated pulses produced smaller MEPs than unpredictable pulses. The predictable computer-generated condition served to test whether the warning cue on its own can attenuate MEPs, in the absence of an intervening response. We found that the warning cue has a small but significant effect on its own, and the response has a larger effect beyond that of the warning cue. This main effect of cue type suggests that the salient internal cues in the lead up to making a response and the high temporal precision afforded by self-generating the stimulation produced greater attenuation in the motor system than signalled computer-generated stimulation using only the warning cue. Moreover, the main effect of contingency confirms that strengthening the causal relationship between actions and cues that trigger TMS pulses produced greater attenuation. This effect of contingency was also consistent across both the predictable conditions (self-generated and signalled computer-generated).

### Experiment 3

Experiment 3 sought to replicate and extend the cue-predictability findings from Experiment 2 by manipulating the temporal precision of an overtly signalled computer-generated TMS pulse. Experiment 2 showed that a warning cue that predicted the arrival of a TMS pulse was sufficient to suppress MEPs. However, the timing of the TMS pulse relative to the onset of the warning cue was yoked to the participant’s response time on the previous trial. This method has the advantage of mimicking individual differences in response timing, but can be affected by outlier response trials. In Experiment 3, we used an on-screen clock to countdown the TMS pulse. This meant that the warning cue (here, the position of the clock-hand aligning with the ‘12 o’clock’ position after completing one rotation) signalled the onset of the TMS pulse with greater temporal precision. Moreover, a countdown procedure rules out the possibility that the reduction in MEPs from the signalled computer-generated condition in Experiment 2 was due to the sudden onset of the warning cue affecting the reflex arc or other reflexive motor cascades separate from prediction mechanisms.

In Experiment 3, the vast majority of trials were clock-TMS trials. For this trial type, most involved a TMS pulse triggered On Time (when the clock-hand reached ‘12’; 8/10 trials), but some were triggered Early (150–450 ms before the clock-hand reached ‘12’; random 1/10 trials) and some were triggered Late (150–450 ms after the clock-hand reached ‘12’; random 1/10 trials). For every 20 clock-TMS trials, we included one Baseline TMS trial where the pulse was triggered during the inter-trial interval (ITI), and one clock trial with no TMS (the clock-hand instead made two complete rotations). The no TMS trial was included so that the Late condition was not entirely predictable if the clock-hand had completed one rotation without triggering a TMS pulse.

**Figure 3.**
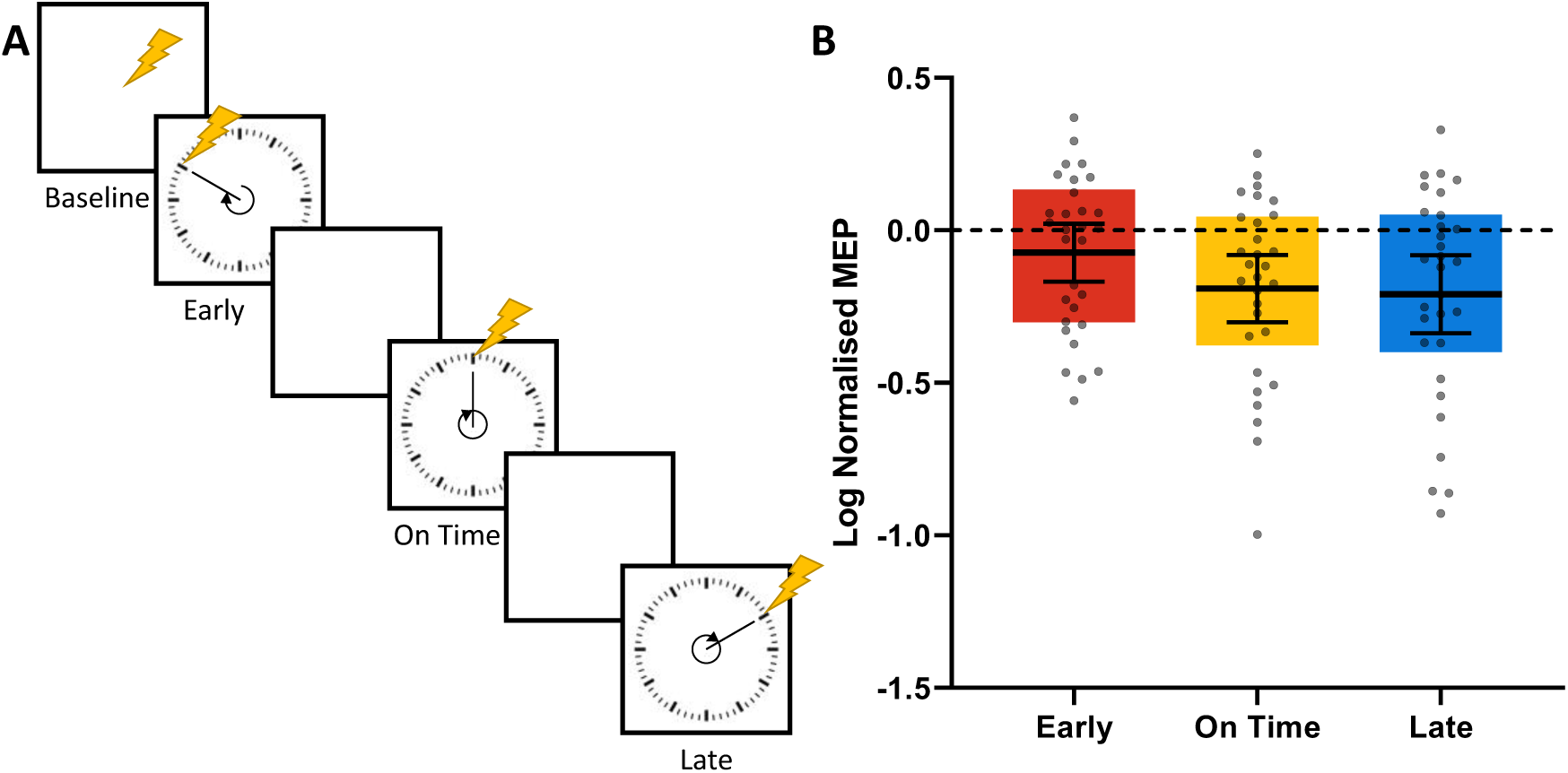
Schematic representation and results of Experiment 3. **A)** Squares represent the screen displayed to participants. A blank screen was used as the default presentation. Participants passively view a clock in which the clock-hand started at the ‘12 o’clock’ position and rotated clockwise. On most trials, TMS was triggered when the clock-hand completed one rotation (On Time condition), which took 1500 ms. On a small proportion of trials, TMS was triggered before (Early condition) or after (Late condition) the clock-hand reached ‘12 o’clock’. TMS was also sometimes randomly triggered during the blank screen (Baseline condition). **B)** Log MEPs normalised to the Baseline condition for each participant. Black bars represent means, and error bars represent 95% CI. Dots represent individual data points, and filled colour boxes represent interquartile ranges. Dashed line represents baseline of log normalisation.

MEPs from the three experimental conditions were normalised to the Baseline condition delivering the unsignalled, unpredictable computer-generated TMS pulses. Stimulation of motor cortex by On Time TMS pulses showed significantly reduced MEPs from baseline (one-sample t-test H_0_ = 0: *t*(29) = 3.53, *p* = 0.001, *d* = 0.64). MEPs generated by Early TMS pulses were not significantly different from baseline (one-sample t-test H_0_ = 0: *t*(29) = 1.56, *p* = 0.126, *d* = 0.29). Interestingly, stimulation by Late TMS pulses showed a pattern more similar to the On Time TMS pulses than the Early pulses, having significantly reduced MEPs from baseline (one-sample t-test H_0_ = 0: *t*(29) = 3.36, *p* = 0.002, *d* = 0.61). This pattern of results was confirmed by a set of orthogonal contrasts showing that normalised MEPs for the Early condition were significantly different from the On Time and Late conditions (*F*(1,29) = 15.45, *p* < 0.001, *η*_*p*_^2^ = 0.35), which did not differ from each other (*F*(1,29) = 0.40, *p* = 0.534, *η*_*p*_^2^ = 0.01).

These results replicate the cued-prediction findings from Experiment 2, showing that, relative to an unsignalled or unpredictable TMS pulse, prediction of a TMS pulse by a reliable warning cue is sufficient to suppress motor system excitability. That is, being able to predict the onset of an externally triggered TMS pulse causes similar, albeit weaker, attenuation to that produced by a self-initiated TMS pulse. However, we found that while Early TMS pulses produced MEPs similar to Baseline TMS stimulation, Late TMS pulses produced MEPs similar to On Time stimulation. Despite Early and Late trials occurring equally infrequently, one explanation for this finding is that Late TMS pulses can largely be predicted by the *absence* of the preceding pulses (Early or On Time). Although there were some trials in which no TMS pulse occurred, if the trial survived one rotation without a TMS pulse, the probability of a Late TMS pulse triggering (p(TMS|after one rotation) = 0.666) was greater than the probability of an Early TMS pulse triggering at the beginning of the trial (p(TMS|before one rotation) = 0.095). We have also conducted another temporal prediction experiment using the same clock presentation but without jittering the time window of the Early and Late conditions, which occurred a fixed 200 ms before or after the On Time condition. This version of the experiment produced the same pattern of results, suggesting that the Experiment 3 findings are also observable without the temporal jitter of the Early and Late conditions. The findings are reported in the Supplementary Materials.

## Discussion

The current series of experiments examined whether attenuation effects arising from prediction mechanisms are present in neural systems beyond the sensory and reward systems, and specifically, in response to stimulation events with which participants have little to no prior experience. We found reliable evidence that predictable or expected TMS pulses triggered over the primary motor cortex resulted in smaller MEPs than unpredictable or surprising TMS pulses. Experiment 1 showed that self-generated TMS pulses produced smaller MEPs (through shifted S-R curves) relative to computer-generated TMS pulses; Experiment 2 showed that signalled TMS pulses (i.e., following a warning cue) produced smaller MEPs than unsignalled TMS pulses; and Experiment 3 showed that temporally predictable TMS pulses produced smaller MEPs than temporally unpredictable TMS pulses. Taken together, the present results provide strong evidence that anticipation or expectation of a TMS pulse causes attenuation of corticospinal excitability throughout the motor system as indexed by MEPs. Here, we demonstrate for the first time that attenuation effects are not specific to the sensory or reward system but are also present in other neural systems and to novel forms of stimulation.

In the sensory domain, it has been argued that an effective sensory system is one in which the organism can easily discriminate, and therefore prioritise, externally-generated sensations from self-generated sensations (Crapse & Sommer, 2008a). Sensory inputs from the external world are less predictable than the recurrent inputs from an organism’s own actions, and may signal novelty or threat in the environment. One way in which the sensory system distinguishes externally- and self-generated sensation is by attenuating the cortical responsiveness to self-generated sensations through a corollary discharge mechanism, which ultimately reduces their perceived salience (Blakemore, Frith, & Wolpert, 1999; Crapse & Sommer, 2008a; Fletcher & Frith, 2009; Kapur, 2003). According to the corollary discharge model, when a motor command is generated, an efference copy of this signal is sent and used to predict the sensory consequences of the action before they are experienced. Once the motor action is performed, the self-generated sensations (sensory reafference) are compared to the predicted sensations (corollary discharge). A match between these signals results in sensory attenuation.

Corollary discharge can be easily reframed in terms of predictive coding. A match between the efference and reafference signals constitutes a correct prediction and thus results in sensory attenuation, while a mismatch between these signals constitutes prediction error and does not result in sensory attenuation. Thinking of corollary discharge in this way is useful for explaining existing sensory attenuation effects in the literature caused by externally-generated sensations that are predictably followed by a warning cue (e.g., Ford et al., 2007). Typically, externally-generated sensations are relatively salient, but the warning cue allows an efference signal to be sent in anticipation of the upcoming sensation. Thus, externally-generated sensations that are signalled become attenuated relative to externally-generated sensations that are unsignalled. Similarly, this predictive coding framework is consistent with findings in which self-generated sensations do not produce sensory attenuation if there is a delay between the motor action of pressing a button to trigger a tone and when the sound plays (e.g., Aliu, Houde, & Nagarajan, 2009; Bäss, Jocobsen, & Schröger, 2008). This is due to the expectation that pressing a button has immediate consequences, and a delayed tone violates this prediction. Critically, if participants receive training to expect the sound to play a short time after pressing the button, then the sensory attenuation effect re-emerges (Elijah et al., 2018). That is, participants can be trained to modify their predictions about the cause and effect relationship between the button press and the arrival of the tone.

Sensory attenuation effects have clear functional benefits that could drive their development on both ontogenetic and phylogenetic timescales. However, our results demonstrate that attenuation effects based on prediction can be observed in non-sensory and non-motivational neural systems. The emergence of such “neural attenuation” effects in the motor system, from a type of direct stimulation with which the participants would have very little or no prior experience, is surprising and reveals commonalities in the general organising principles of neural circuits across the brain. These findings suggest that predictive coding may be a domain-general language of the brain, present across many neural systems and potentially more widespread than initially thought. Extending this idea, the corollary discharge mechanisms responsible for numerous sensory attenuation effects observed across species including, fish, birds, and monkeys (Crapse & Sommer, 2008a) may be a specific form of the predictive coding architecture developed by the sensory system.

The attenuation findings in the motor system also have important practical implications for testing and designing TMS experiments in Cognitive Neuroscience as well as using TMS for therapeutic purposes in clinical setttings. The typical procedure for determining rMTs usually requires an experimenter to find the lowest percentage of stimulator output that produces 5 out of 10 trials with MEPs at least as large as 50 µV. However, the way in which TMS pulses are triggered during this set-up procedure can affect how predictable the participants find the stimulation and therefore the size of the resulting MEP. For example, if the participant is self-triggering pulses or if the experimenter triggers pulses at a regular interval then this will produce smaller MEPs. This presents a practical issue for TMS researchers because it means that the stimulator value may not be set at an ideal position on the dose-response function for peak sensitivity to experimental manipulations. Of greater concern, however, is that it creates issues of variability between labs conducting similar experiments or clinical trials but setting different rMTs based on variations in the procedure.

A second methodological consideration revealed by our findings is the importance of selecting an appropriate baseline condition when designing experiments. Baseline conditions are useful for normalising data and reducing large individual differences in the absolute values of MEP amplitudes. However, using a baseline condition in which the TMS is triggered unpredictably to normalise an experimental condition in which the TMS is predictable (e.g., timed regularly relative to a stimulus onset) will at best weaken any excitatory effects and at worst lead to incorrect interpretations of a suppression effect due to the attenuated MEP relative to baseline. Therefore, it is important to design TMS experiments in which baseline trials are matched in predictability to the experimental trials.

In summary, we have shown that predictable stimulation of the motor cortex leads to attenuation of the resulting MEPs relative to unpredictable stimulation. The results reveal that neural attenuation effects arising from prediction are present in the brain beyond the sensory and motivational systems and to novel forms of stimulation. The findings also have significant theoretical and practical implications, offering new insights into the generality of predictive coding throughout the brain, and emphasising the importance of standardising rMT procedures as well as designing experiments with appropriate baseline conditions.

## Method and Materials

### Participants

All participants completed a TMS safety screening and provided their informed consent before starting the experiment or any set-up procedures. All experimental protocols were approved by the Human Research Ethics Committee of The University of Sydney.

#### Experiment 1

The experiment was conducted on 10 right-handed participants (1 female; mean age = 29.80, SD = 5.45), including three of the authors. Since this experiment involved setting the TMS machine to very high percentages of the maximum stimulator output (MSO), this restricted our sample size and all the participants were known to the authors.

#### Experiment 2

We aimed to have 24 participants included in the study (12 participants in the hand and 12 participants in the foot condition). The experiment was conducted on 25 right-handed participants from The University of Sydney first-year Psychology testing pool. The data were analysed on 24 participants (14 females; mean age = 24.63, SD = 6.02) after one participant withdrew due to the level of stimulation required to elicit reliable MEPs.

#### Experiment 3

We aimed to have at least 24 participants included in the study after exclusions. We set out to recruit 30 participants from The University of Sydney first-year Psychology testing pool, and 33 participants registered. The experiment was conducted on all 33 participants. The data were analysed on 30 participants (1 left-handed; 22 females; mean age = 22.20, SD = 4.28) after a technical error did not save one participant’s data and two participants were excluded during the data processing stage (detailed below).

### Electromyography and TMS

TMS was administered using a Magstim 2002 with a 70-mm figure-eight coil (Magstim, Whitland, UK). Pulses were delivered to the left primary motor cortex (M1) with the coil positioned in a posterior–anterior configuration, with the handle oriented approximately 45° from midline. Surface electromyography (EMG) traces were recorded from the FDI muscle of the right hand, using a pair of Ag/AgCl electrodes placed in a belly–tendon arrangement over the muscle along with a ground electrode placed over the ulnar styloid process of the wrist. Data from 200 ms pre-stimulation to 100 ms post-stimulation were collected via a PowerLab 26T DAQ device (ADInstruments, Bella Vista, NSW, Australia). The analogue EMG signal was digitized (sampling rate: 4 kHz; bandpass filter: 0.5 Hz to 2 kHz; mains filter: 50 Hz) and stored on a computer using LabChart software (Version 8, ADInstruments) for offline analysis. Timing between TMS and EMG was synchronized through TTL signals sent from the test computer to the Magstim unit and PowerLab.

The location of the motor cortex “hotspot” was determined starting from a spot 5 cm lateral and 1 cm anterior to Cz and moving the coil until the maximal MEP was elicited in the FDI muscle. A fitted cap with the 10–20 EEG locations marked was worn by participants to aid in this process. Once the hotspot was located, an adjustable forehead and chin rest was used to minimise any subsequent movement and the coil was locked in position with the aid of a mechanical arm (Manfrotto, Cassola, Italy).

#### Experiments 2 & 3

Resting motor threshold (rMT) was defined as the lowest stimulation intensity capable of inducing MEPs with a minimum of 50 μV peak-to-peak amplitude in 5 of 10 consecutive pulses (Rossini et al., 2015). Pulse intensity during the experiment was then set at 120% of rMT. For Experiment 2, the mean rMT across participants was 55.29% MSO (SD = 10.29%), and for Experiment 3, the mean rMT was 44.43% MSO (SD = 8.41%).

### Apparatus and Stimuli

All experiments were run on a Windows 7 PC using PsychoPy2 (Experiments 1 and 2) or MATLAB software (Experiment 3) to control stimulus presentation. Stimuli were displayed on a 24-inch ASUS monitor (1920 × 1080 resolution, 60 Hz refresh rate) at a viewing distance of ∼57cm. The text stimuli displayed onscreen subtended approximately 2° of visual angle.

### Experiment Procedure

#### Experiment 1

The default display throughout Experiment 1 was a fixation cross presented in the centre of the screen. TMS pulses could be triggered in three different ways: 1) the participant pressing the spacebar with their left thumb in response to the word “Thumb*”* presented on the screen (Thumb condition); 2) the participant pressing the spacebar on a second keyboard on the floor with their left foot in response to the word “Foot” presented on the screen (Foot condition); or 3) directly via the experiment script with no prior cue or warning to the participant or change to the display (Computer condition). For the Foot and Thumb conditions, the word prompt remained on display until the spacebar was pressed. The word disappeared and was replaced with a fixation cross as soon as the spacebar was pressed. Participants were informed at the beginning of the experiment to respond at their own comfortable pace, and to prioritise accuracy of the effector used to make the response over speed. The order of triggering type or conditions within each block was pseudo-randomised such that no condition occurred more than twice in a row.

S-R curves for each condition were collected using a rapid method based on Mathias et al. (2014). Data were acquired for each curve by delivering a single TMS pulse at each intensity value from 20 to 100% of MSO; a total of 81 pulses per SR curve. The experiment consistent of three blocks and the intensity of pulses was controlled such that they were identical for each condition as well as being evenly spread across the full range of possible values. For example, one block used the intensities 20 – 23 – 26 – … 98% MSO for all conditions, the second block used 21 – 24 – 27 – … 99% MSO, and the third block used 22 – 25 – 28 – … 100%. The actual assignment of these intensity ranges to blocks was randomised across participants, and within each block the intensities were delivered in a randomised order. Each block contained 81 TMS pulses (27 per condition), and the minimum interval between pulses varied between 6 and 9 seconds. There was a self-timed break between each block and a total of 243 trials across the 60 min experiment (including briefing, set-up, and debrief). If a particular interval was too short to allow adequate time for the Magstim to change the intensity from the previous pulse, it was extended to the required duration. Control of the Magstim stimulator was achieved using the MagPy toolbox (McNair, 2017). This Python toolbox provided full programmatic control to the experiment script over changing stimulator intensity, arming/disarming of the stimulator, and firing of the TMS pulse.

A BrainSight neuronavigation system (Rogue Research, Montreal, QC, Canada) was used to ensure the coil remained in position over the identified motor hotspot when delivering TMS pulses. In some cases, a participant’s own structural MRI was used, while the MNI152 brain template was used for the remaining participants. Coregistration of the participant’s head with the MRI image was achieved by aligning three fiduciary markers at the nasion and left and right preauricular points. Once the coil was locked into place over the hotspot, this target position was tracked via the BrainSight software. Trials in which the coil moved more than 5 mm from the hotspot were excluded from subsequent analysis.

#### Experiment 2

TMS pulses could be triggered in three different ways: 1) the participant pressing the spacebar with their left thumb (n = 12) or their left foot (n = 12) in response to one shape appearing in the centre of the screen (e.g., “**+**”; Self-generated condition); 2) automatically following the presentation of a second shape appearing in the centre of the screen (e.g., “**o**”; Warning cue condition); or 3) during on the blank screen between two ITI periods (Unpredictable condition). At the start of the experiment, participants were instructed which shape required a key press and which shape did not, and the stimuli were counterbalanced between participants. There was no emphasis on the speed of responding and participants were explicitly informed to press the key whenever they were ready. The TMS pulse was triggered immediately on the press of the spacebar for the Self-generated condition, or after a short delay following the Warning cue and Unpredictable conditions. For Warning cue trials, the delay between the onset of the stimulus and the TMS pulse was equal to the response time from the most recent Self-generated trial. Similarly, for Baseline trials, the TMS pulse was triggered after a variable ITI following a short delay equal to the response time from the most recent Self-generated trial. If there had not been a Self-generated trial before the first Warning cue or Baseline trial, the delay was 300 ms. There were no stimulus presentation changes during the ITI and Baseline trials, with the screen remaining blank until the next Self-generated or Warning cue trial. On Self-generated and Warning cue trials the stimulus remained on the screen for 1500 ms after the delivery of the TMS pulse. Trials were pseudo-randomised such that no condition occurred more than four times in a row and the ITI period jittered randomly between 2000 and 3000 ms.

The experiment was split into two phases, a low and high contingency phase. In the low contingency phase, participants received a TMS pulse on a random 1 in 4 trials of the same condition. There were 360 trials in the low contingency phase, 120 trials of each condition, and 30 trials (1:4) of which triggered a TMS pulse. In the high contingency phase, participants received a TMS pulse on every trial. There were 90 trials in the high contingency phase, 30 trials of each condition. A self-timed break was scheduled after each block and between the two phases; participants could choose to continue the experiment or stop and have the TMS coil removed and could move away from the forehead and chin rest for a stretch. One block included 36 trials in the low contingency phase and 30 trials in the high contingency phase. The total experimental session including briefing, set-up, testing, and debrief was approximately 60 minutes.

#### Experiment 3

At the start of every trial, a clock appeared in the centre of the screen with the clock-hand starting at the ‘12 o’clock’ position and rotating clockwise. On 8/10 trials, the TMS pulse was triggered when the clock-hand made one complete rotation back to the ‘12 o’clock’ position (On Time condition), which took 1500 ms. On 1/10 trials the TMS pulse was triggered 150-450 ms (approximately between the 8 – 11 o’clock positions) before completing one complete rotation (Early condition). On 1/10 trials the TMS pulse was triggered 150-450 ms (approximately between the 1 – 4 o’clock positions) after completing one complete rotation (Late condition). For every 20 clock-TMS trials, there was 1 clock trial in which the clock-hand made two complete rotations without delivering a TMS pulse (no TMS trial), and 1 trial in which the clock did not appear while TMS was triggered on a blank screen between two ITI periods (Baseline condition). The ITI period jittered randomly between 2500 and 4000 ms. The experiment was divided into six blocks separated by a self-timed break for participants to have the TMS coil removed and stretch. Each block included 66 trials for a total of 396 trials across the 60 min experiment (including briefing, set-up, and debrief).

## Data Analysis

Pre-processing of MEP data was conducted using custom software written in Python. Trials with background EMG larger than 50 μV peak-to-peak amplitude prior to the TMS pulse were excluded. In addition, trials with EMG activity greater than 100 μV RMS over the entire 200 ms pre-stimulation period were also removed from analysis. MEP amplitudes were then measured from the remaining trials using peak-to-peak difference values.

### Experiment 1

The relationship between TMS intensity (*X*) and MEP amplitude (*Y*) was modelled in MATLAB (2018a; The Mathworks Inc., Natick, MA, USA) using a four-parameter Sigmoidal-Boltzmann function:

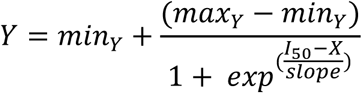

Where *min*_*Y*_ and *max*_*Y*_ are the minimum and maximum MEP amplitude asymptotes, respectively, *I*_*50*_ is the stimulator intensity at the inflection point halfway between *min* and *max*, and *slope* is the steepness of the curve. A least absolute residuals method was used to achieve a robust fit of the data, by giving equal weighting to all data points, and to reduce the effect of extreme values. Largely following Mathias et al. (2014), initial parameters were set as follows: *max*_*Y*_ = max(Y); *I*_*50*_ = 60%; and *slope* = 5; while *min*_*Y*_ was fixed at 0. The *max*_*Y*_ and *slope* parameters were constrained to be greater than 0, while *I*_*50*_ was constrained to fall between 25 and 95% MSO.

### Experiments 2 & 3

Since the BrainSight neuronavigation system was not used in Experiments 2 and 3, any MEP less than 50 µV was treated as a mistrigger that occurred off the hotspot location and was excluded from the analysis. Additionally, two participants were excluded from Experiment 3; one participant did not produce reliable MEPs at 120% of rMT (more than half of the TMS trials were < 100 µV), and another participant had mean MEP amplitudes more than 3 SD below the group mean across 2 of the 3 conditions.

The MEP data were log-normalised at a participant level to remove large individual differences in the absolute MEP values. Normalising MEPs is commonly adopted in the TMS literature, however, raw normalisation (e.g., 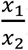) centers the midpoint around a value of 1 and creates a positively skewed distribution that favors “excitation” (*x*_1_ > *x*_2_; 1 → ∞) over “suppression” (*x*_2_ > *x*_1_; 0 → 1). Some researchers have attempted to recentered the mid-point around a value of 0 using a linear transformation (e.g.,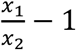), but this does not change the skew of the distribution or bias for excitation over suppression effects. Log-transforming the normalised values centers the mid-point around 0 and makes the distribution symmetrical, thereby allowing excitation and suppression effects to span the same range of values from 0 → +∞ and 0 → -∞, respectively. We have adopted log-normalisations in our previous work (Tran et al., 2019; Tran et al., online ahead of print). Average raw MEPs are also presented in the Supplementary Materials.

## Acknowledgements

The authors thank Dr Nahian Chowdhury for his assistance with data collection. The research was supported by the Australian Research Council’s Discovery Projects funding scheme (project number DP190100410).

