## Supplementary Materials for "Towards domain-general predictive coding: Expected TMS excites the motor system less effectively than unexpected stimulation"

#### Parameter values

**Table S1.** Mean (SEM) goodness of fit ($R^2$ adjusted) and parameter values for sigmoid functions from Experiment 1. The maximum parameter value was normalised to the Computer condition.

| | Goodness of fit | Maximum | Slope | Inflection |
|---|---|---|---|---|
| Computer | 93.154 (3.451) | 1.000 (0.000) | 4.759 (0.647) | 54.866 (2.429) |
| Foot | 93.235 (3.417) | 0.958 (0.048) | 5.315 (0.735) | 58.331 (2.960) |
| Thumb | 93.617 (3.095) | 0.863 (0.066) | 4.888 (0.572) | 56.389 (2.782) |

#### Behavioural data

Response time (RT) data were not recorded for Experiment 1. In Experiment 2 the response time from the previous Self-generated trial was used to yoke the TMS delay period for the subsequent Warning cue trial. RTs more than 3 standard deviations above a participant's mean were excluded, this removed an average of 2.46 (SD = 1.25) trials per participant. The mean RT for participants who self-generated a TMS pulse with their left foot (n = 12) was 667.39 (SD = 165.36) ms and for those who self-generated a TMS pulse with their left thumb (n = 12) was 551.11 (SD = 91.76) ms. As an indication of within-participant RT variability across trials, the mean standard deviation calculated from the self-generated trials for each participants was 131.61 (SD = 73.26) ms.

### Raw MEP data

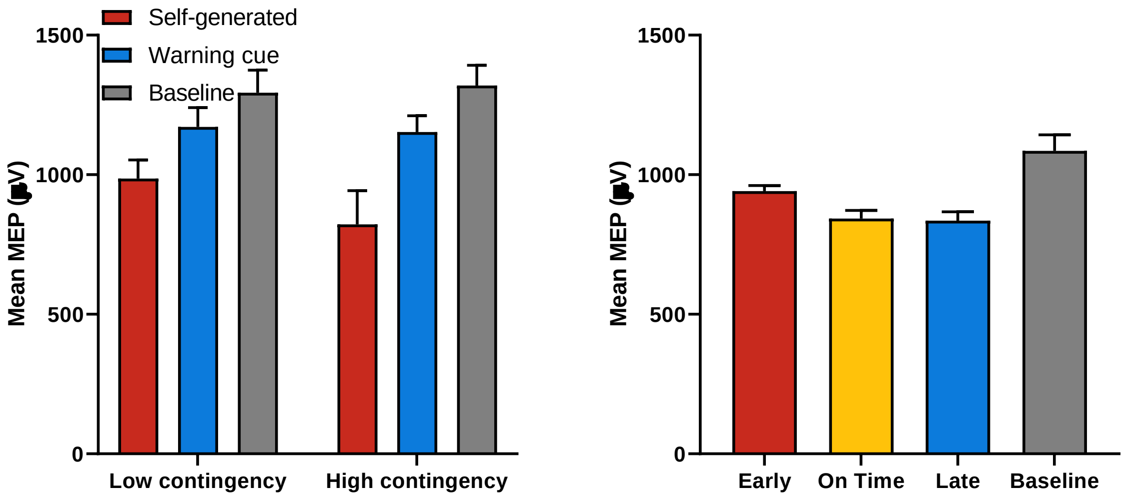


**Figure S1.** Mean raw MEPs for Experiment 2 (left) and Experiment 3 (right). Error bars represent within-participant standard errors (Morey, 2008).

### RAW RMS data

We also analysed the EMG trace before the TMS pulse to check that the prediction effects on MEP are independent of any baseline EMG activity from the participants tensing or bracing for the delivery of upcoming predictable pulse, which would manifest as increased background EMG noise during the foreperiod. To do this, we calculated the root mean squared (RMS) EMG activity from -50 to -3 ms in the lead up to the TMS pulse.

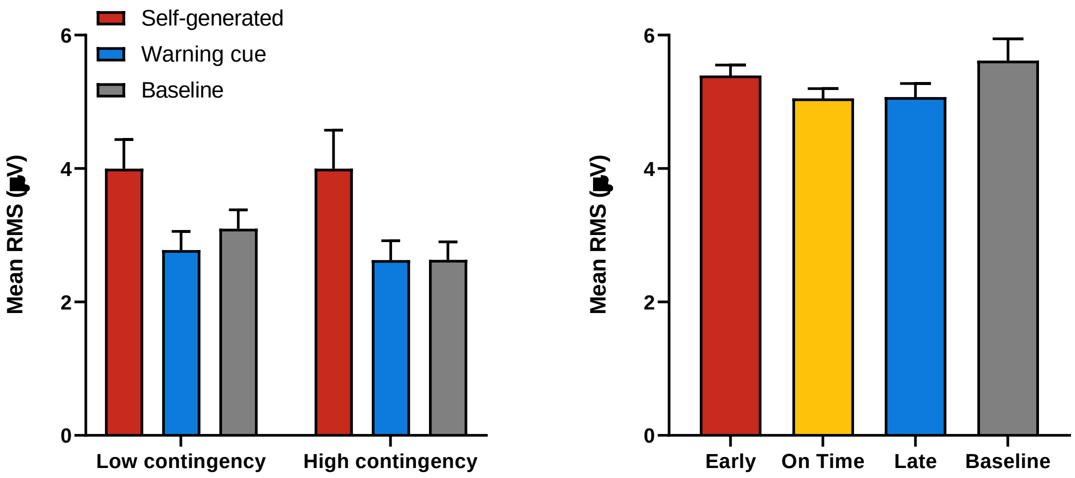


**Figure S2.** Mean raw RMS for Experiment 2 (left) and Experiment 3 (right). RMS was calculated from -50 to -3 ms before the TMS pulse. Error bars represent within-participant standard errors (Morey, 2008).

For Experiment 2, there was a main effect of trial type [self-generated, warning cue, baseline] on RMS ($F(1,46) = 3.43$, $p < 0.041$, $\eta_p^2 = 0.13$), and no effect of contingency [low, high] ($F(1,46) = 0.90$, $p = 0.353$) or interaction ($F(1,46) = 1.14$, $p = 0.330$). This appears to reflect higher RMS in the condition that requires action (self-generated) compared to the other two, rather than a difference between the predictable and unpredictable conditions. A set of orthogonal contrasts comparing the predictable [self-generated, warning cue] and unpredictable conditions [baseline] ($F(1,23) = 2.14$, $p = 0.157$), and the predictable conditions against each other [self-generated v warning cue] ($F(1,23) = 3.87$, $p = 0.061$) were both non-significant. These results suggest the effect of trial type on RMS is driven by the requirement to respond, rather than the anticipation of a TMS pulse as RMS is higher for self-generated trials compared to warning cue and baseline trials.

For Experiment 3, there was no overall effect of trial type ($F(3,87) = 1.51$, $p = 0.217$). A set of orthogonal contrasts comparing the predictable [on time, late] and unpredictable [early, baseline] conditions ($F(1,29) = 3.44$, $p = 0.074$), the predictable conditions against each other [on time v late] ($F(1,29) = 0.35$, $p = 0.558$), and the unpredictable conditions against each other [early v baseline] ($F(1,29) = 0.01$, $p = 0.921$) were all non-significant. The contrast comparing the predictable and non-predictable conditions was marginally significant, but is in the *opposite* direction to that predicted by the participant tensing or bracing in anticipation of the pulse. Instead, RMS was slightly higher for the unpredictable conditions.

Together, the background EMG data from Experiments 2 and 3 suggest that anticipation from the TMS pulse as measured by RMS is unlikely to account of the prediction effects on MEP amplitudes.

**Supplementary Experiment 1**

We aimed to have at least 24 participants included in the study after exclusions. We set out to recruit 30 participants from The University of Sydney first-year Psychology testing pool, and 30 participants registered. The experiment was conducted on all 30 participants. The data were analysed on 28 participants after two participants were excluded for not completing the study.

Supplementary Experiment 1 included three conditions, an On Time condition (TMS pulse generated when the clock-hand reached ‘12 o’clock’; 8:10 trials), and two conditions where a TMS pulse was either triggered Early (200 ms before the clock-hand reached ‘12’; random 1:10 trials) or Late (200 ms after the clock-hand reached ‘12’; random 1:10 trials). MEPs from the On Time and Late conditions were normalised with respect to the Early condition. Stimulation of the primary motor cortex by On Time TMS pulses showed the expected reduction in MEP amplitude relative to the Early condition (one-sample t-test $H_0 = 0$: $t(27) = 2.60$, $p = 0.015$, $d = 0.49$). Stimulation by Late TMS pulses also showed significantly reduced MEPs relative to the Early condition (one-sample t-test $H_0 = 0$: $t(27) = 2.26$, $p = 0.032$, $d = 0.43$), and normalised MEPs from the On Time condition did not differ significantly to those in the Late condition ($t(27) = 0.88$, $p = 0.388$, $d = 0.17$). These results replicate the findings from Experiments 2 and 3 in the main article showing that predictable stimulation of On Time pulses suppressed MEPs relative to unpredictable stimulation of Early pulses. They also replicate the results from the Late condition producing an attenuation pattern similar to the On Time condition, despite occurring just a randomly and infrequently as the Early condition. As previously explained, Late TMS pulses can be predicted by the *absence* of both Early and Predictable pulses. That is, if a TMS pulse has not been delivered by the time the clock hand has passed the ‘12 o’clock’ position then participants can learn to expect that there would be an upcoming Late pulse. Although the delay was only 200 ms, it is likely enough time for participants to adjust their predictions.

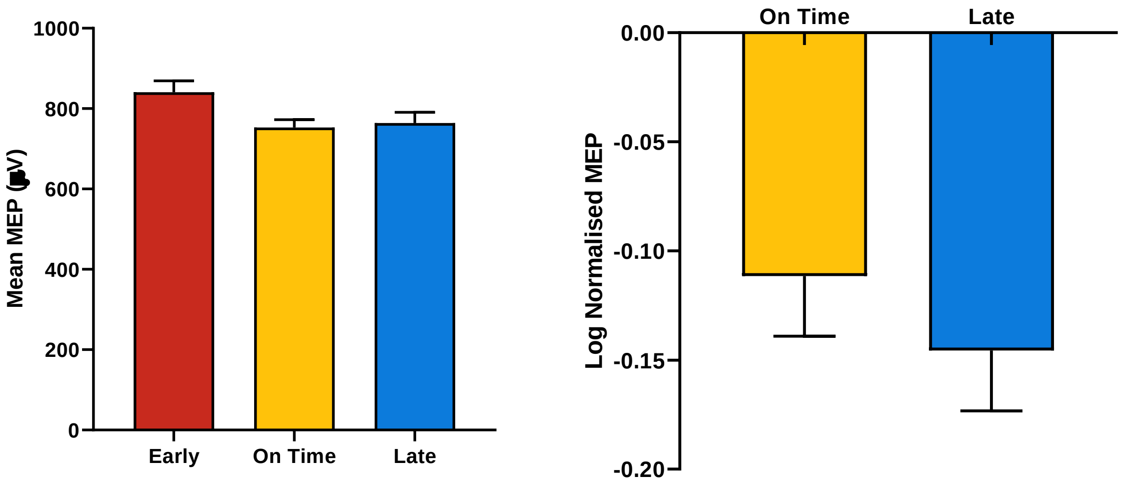


**Figure S3.** Mean raw MEPs (left) and log MEPs normalised to the Early condition for each participant (right). Error bars represent within-participant standard errors (Morey, 2008).
